# Behavioral response to visual motion impacts population coding in the mouse visual thalamus

**DOI:** 10.1101/382671

**Authors:** Karolina Socha, Matt Whiteway, Daniel A. Butts, Vincent Bonin

## Abstract

Visual motion is a ubiquitous component of animals’ sensory experience and its encoding is critical for navigation and movement. Yet its impact on behavior and neural coding is not well understood. Combining pupillometry with cellular calcium imaging measurements of thalamocortical axons in awake behaving mice, we examined the impact of arousal and behavioral state on encoding of visual motion in the visual thalamus. We discovered that back-to-front visual motions elicits a robust behavioral response that shapes tunings of visual thalamic responses. Consistent with an arousal mechanism, the effects were pronounced during stillness and weak or absent during locomotor activity and under anesthesia. The impact on neuronal tuning was specific, biasing population response patterns in favor of back-to-front motion. The potent influence of visual motion on behavioral state dynamically affect sensory coding at early visual processing stages. Further research is required to reveal the circuitry and function of this novel mechanism.

## Introduction

Animal behavior involves a continuous interplay between encoding of sensory inputs and updating of internal states enabling critical functions and flexible and adaptive responses. This interplay is particularly striking in the neuronal processing of visual motion which is critical for navigation (Warren and Hannon, 1988; Srinivasan et al., 1999; Mertes et al., 2014; Shiozaki and Kazama, 2017), eye movements (Warren and Hannon, 1990), whole body movements (Lappe et al., 1999) and can trigger innate behaviors (Yilmaz and Meister, 2013; De Franceschi et al., 2016; Salay et al., 2018; Shang et al., 2018).

Encoding of visual motion begins with direction selective responses which are tuned to stimuli moving in particular directions. While direction selective responses are seen throughout the brain (Hubel and Wiesel, 1962; Barlow and Hill, 1963; Oyster and Barlow, 1967; Livingstone, 1998; Inayat et al., 2015), a fraction in mice can be traced back to activity of direction selective retinal ganglion cells (Cruz-Martín et al., 2014; Hillier et al., 2017). These cells are preferentially tuned to cardinal directions of motion, forming a population code for global motion (Briggman et al., 2011; Kay et al., 2011; Dhande et al., 2013; Yonehara et al., 2013; Sabbah et al., 2017). These cells send their outputs to neurons in the dorsal lateral geniculate nucleus (dLGN) neurons, which distribute visual motion signals to distinct layers of the primary visual cortex (Cruz-Martín et al., 2014; Kondo and Ohki, 2015; Roth et al., 2015; Sun et al., 2015).

However, there is considerable disagreement in the degree to which different directions of motion are represented in areas downstream of the retina. Similar to what is observed in the retina, a study of dLGN inputs to V1 in head-fixed mice reported a nearly equal numbers of V1 inputs tuned to each of the four cardinal directions (Kondo and Ohki, 2015). Another study reported instead a pronounced over-representation of inputs tuned to stimuli moving in the general temporal-to-nasal direction (Sun et al., 2015). Considerable diversity in response tuning has also been observed in the mouse cortex (Andermann et al., 2011; Rochefort et al., 2011; Sun et al., 2015; Hillier et al., 2017; Marques et al., 2018). The inconsistency in the dLGN data could reflect genuine differences in visual computations in the dLGN. Alternatively, it could reflect an unknown mechanism that shapes dLGN responses to visual motion stimuli.

A possible explanation are behavioral modulations which are known to influence early visual responses (Swadlow and Weyand, 1987; McAdams, 2005; McAlonan et al., 2008; Saalmann and Kastner, 2009; Niell and Stryker, 2010; Keller et al., 2012; Ayaz et al., 2013; Bennett et al., 2013; Saleem et al., 2013; Polack et al., 2013; Erisken et al., 2014; Hei et al., 2014; Roth et al., 2015; Vinck et al., 2015; Busse et al., 2017; Wang and Krauzlis, 2018). These modulations have been shown to affect the visual responsiveness and the tunings of dLGN neurons and therefore could potentially explain the differences in tunings observed across studies. Interestingly, the studies by (Kondo and Ohki, 2015) and (Sun et al., 2015) were conducted in anesthetized and awake animals which suggests behavioral modulations as a potential explanation. Furthermore, visual motion stimuli have been described to innate behaviors (Yilmaz and Meister, 2013; De Franceschi et al., 2016; Salay et al., 2018; Shang et al., 2018). Such behaviors likely engage neuromodulatory mechanisms that may impact neuronal encoding of visual motion stimuli. While studied extensively in insects (Borst and Haag, 2002; Egelhaaf et al., 2002; Boeddeker et al., 2005; Maimon et al., 2010; Jung et al., 2011; Reiser and Dickinson, 2013; Borst, 2014), the impact of visual motion on behavioral state and sensory encoding has not been systematically investigated in mice.

In this study, we investigated in head-fixed locomoting mice the impact of arousal and behavioral state on encoding of visual motion in the visual thalamus. We discovered that visual motion stimuli elicit a specific arousal response. This arousal response strongly modulate dLGN response to visual stimuli biasing population measures of direction selectivity and preferences. The results demonstrate that visual motion can exert a potent influence on the behavioral state impacting sensory coding at early stages of visual processing. They reconcile differences in population tuning reported in past studies and suggest that dLGN neurons faithfully carry retinal representations of visual motion from the retina to the cortex.

## Results

To study the encoding of visual motion in the visual thalamus, we combined cellular two-photon imaging with pupillometric measurements in head-fixed mice (Figure 1). Mice were imaged on a treadmill while receiving visual motion stimuli and performing voluntary locomotor behavior (Figure 1A). We measured both the behavioral and neuronal responses to luminance-calibrated sinusoidal drifting gratings of fixed spatial and temporal frequency (0.08 cpd, 4 Hz) and varying orientation and direction (Figure 1B), as used in studies of motion direction selectivity (Cruz-Martín et al., 2014; Kondo and Ohki, 2015; Roth et al., 2015; Sun et al., 2015). These stimuli were presented to the right visual field (120 deg by 80 deg, width by height) in 5-sec intervals interleaved with epochs of static grey screen.

**Figure 1.**
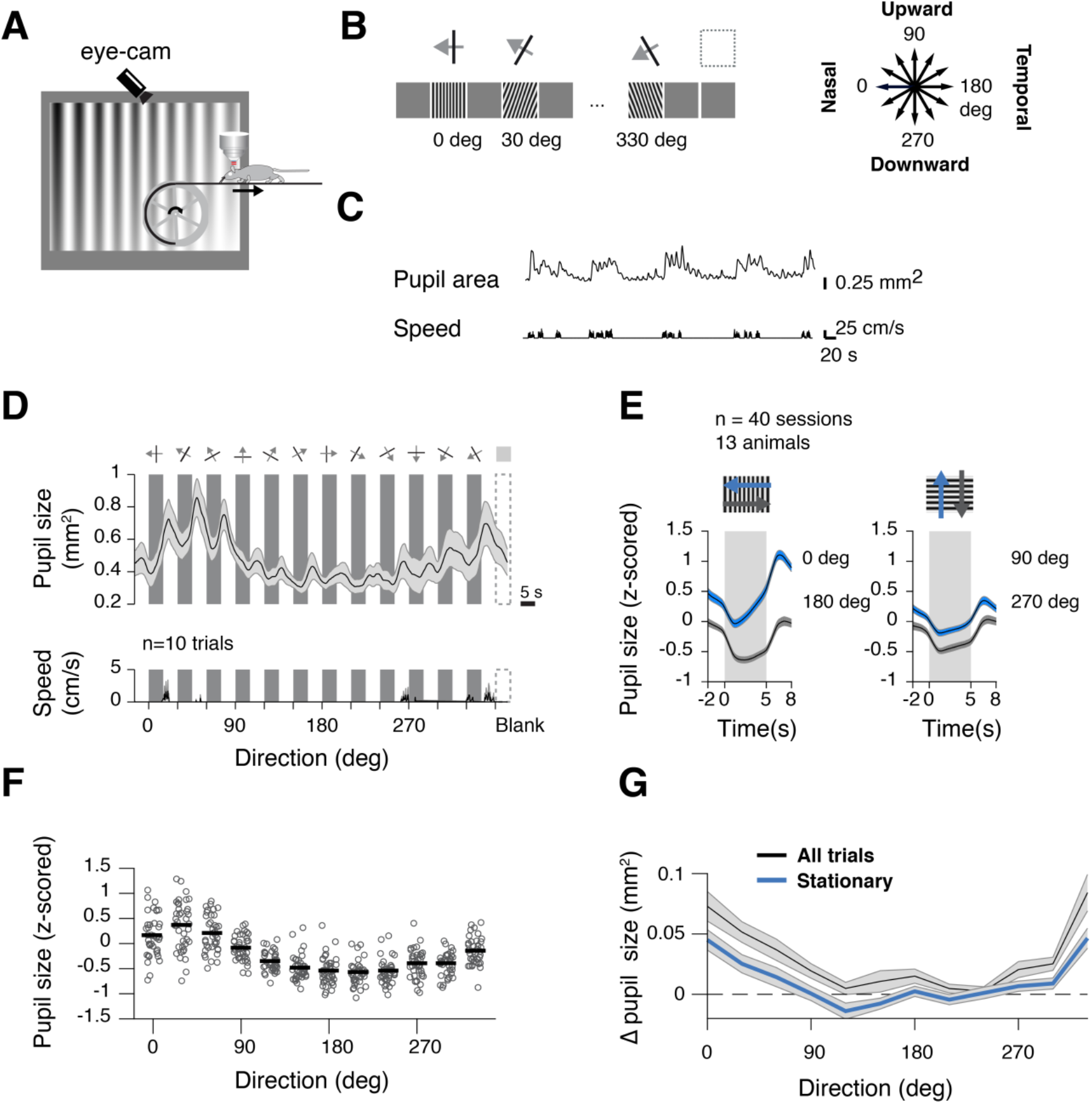
Temporal-to-nasal visual motion increases pupil size in head-fixed mice. (A) Illustration of the voluntary locomotor assay. Mice are head-fixed on a linear treadmill and allowed to run while pupil size and locomotion speed are measured. (B) Diagram of visual stimulus sequence. Sinusoidal drifting gratings (120 deg by 80 deg, width by height) moving in one of 12 directions were presented sequentially in 5-s intervals interleaved with equiluminant gray screen. (C) Example pupil size and locomotion speed measurements. (D) Example pupil response to the stimuli. (mean±SD); Shaded areas indicate stimulation epochs, white – blank periods; dashed lines – blank stimulus. (E) Normalized pupil responses triggered by the onset of stimulus for the cardinal directions (mean±S.E.M). Shaded area is stimulus duration; colors differentiate opposing directions. Pupil increases in response to back-to-front moving gratings but not the opposite direction. There is no change in pupil response for upward and downward directions. (F) Effect of stimulus direction on pupil size. Grey dots are the average pupil size for each session during 5s of stimulation. Black lines are the mean across all session. (G) Relative change in pupil size (difference between pupil size in the last second of the stimulus and baseline) for gratings in 12 directions; mean±S.E.M. Black – all trials, blue – stationary trials (speed <1 cm/s for at least 95% of stimulus duration). See also Figure S1.

The animals ran occasionally (25±22% of the session; n=40 sessions) allowing comparison of eye movements and neural activity across locomotor states. Consistent with previous work (Erisken et al., 2014; Vinck et al., 2015) pupil size strongly covaried with locomotor activity (Figure 1C), increasing sharply during movement, and decreasing slowly during stillness. These locomotion-related changes in pupil size were described previously and are believed to reflect changes in arousal (Bennett et al., 2013; Reimer et al., 2014; Vinck et al., 2015).

### Temporal-to-nasal visual motion increases pupil size in head fixed mice

We first examined the impact of the visual motion stimuli on arousal and locomotor state (n=13 mice, 40 sessions). Visual motion can elicit behavioral responses (Yilmaz and Meister, 2013; De Franceschi et al., 2016; Salay et al., 2018; Shang et al., 2018) that could potentially modulate visual responses. While drifting gratings stimuli have been used in numerous awake rodent studies (Niell and Stryker, 2010; Andermann et al., 2011; Glickfeld et al., 2013; Sun et al., 2015; Durand et al., 2016), their impact on behavioral state has never been systematically characterized. We related locomotion speed, eye movements, and pupil size, a proxy for arousal, to visual stimulus onset and offset (Bennett et al., 2013; Reimer et al., 2014; Vinck et al., 2015).

We observed strong pupil size responses to the visual motion stimuli on the display (Figure 1D-G). During stimulation epochs, visual motion stimuli led to progressive dilation of the pupil that lasted through the visual stimulation epochs (Figure 1D). These pupil responses were pronounced (Figure 1E) and visible in single trials (Figure 1C, Figure S1A). During grey screen epochs, the pupil also showed a brief increase followed by a slow relaxation towards baseline that depended on the response to the preceding stimulus. The response to the visual motion stimuli showed a strong dependence of direction of motion on the display (Figure 1D, top). The transient response at stimulus offset, in comparison, was relatively weak and unspecific present for all stimuli (Figure 1E, Figure S1B). In this report, we focus solely on the increase in pupil size during visual stimulation epochs.

The pupil responses to the visual stimuli showed a striking bias for direction of motion (Figure 1E). Pronounced increases in pupil size were observed for stimuli drifting in the temporal-to-nasal directions (Figure 1E, left, blue). No responses were observed for stimuli moving in upward, downward and nasal-to-temporal directions (Figure 1E, right, grey). Consistent with a response to back-to-front visual motion, magnitude of the pupil response decreased gradually with gratings’ direction of motion (Figure 1D,F,G, Figure S1B). Critically, these responses occurred without concomitant locomotor activity (Figure 1D bottom, Suppl. S1A). The bias for direction was robust, seen consistently across animals (Figure 1F). While visual stimuli could occasionally trigger or modulate locomotor activity (Figure S1C), the bias for direction was also seen in stationary trials (Figure 1G).

### Pupil size responses to visual motion reflect a vision-only arousal mechanism

The specific pupil responses to visual stimuli could reflect a visual mechanism in which back-to-front motion stimuli arouse the animal eliciting a pupil response. Alternatively, the response could reflect a visuomotor startled response to mismatch between actual visual inputs and those expected from self-motion (Keller et al., 2012).

To distinguish between these possibilities, we examined pupil responses to the visual stimuli as a function of locomotor state and movement speed. If pupil fluctuations reflect a sensorimotor mismatch response, then a shift of the effects towards front-to-back direction should be observed during running. If pupil responses instead reflect a vision-only arousal response to back-to-front visual motion, then the arousal effects of visual motion should decrease in magnitude when animal is engaged in locomotion.

Consistent with the vision-only hypothesis, we found during locomotor activity a dramatic weakening of the pupil response to back-to-front motion as well as no evidence of response to front-to-back visual motion (Figure 2). The effects of visual stimulation on pupil size were strongest during still epochs (Figure 2A) and weakest during locomotion (Figure 2B). Locomotion also largely abolished the bias of pupil size for back-to-front motion (Figure 2C,D).

**Figure 2.**
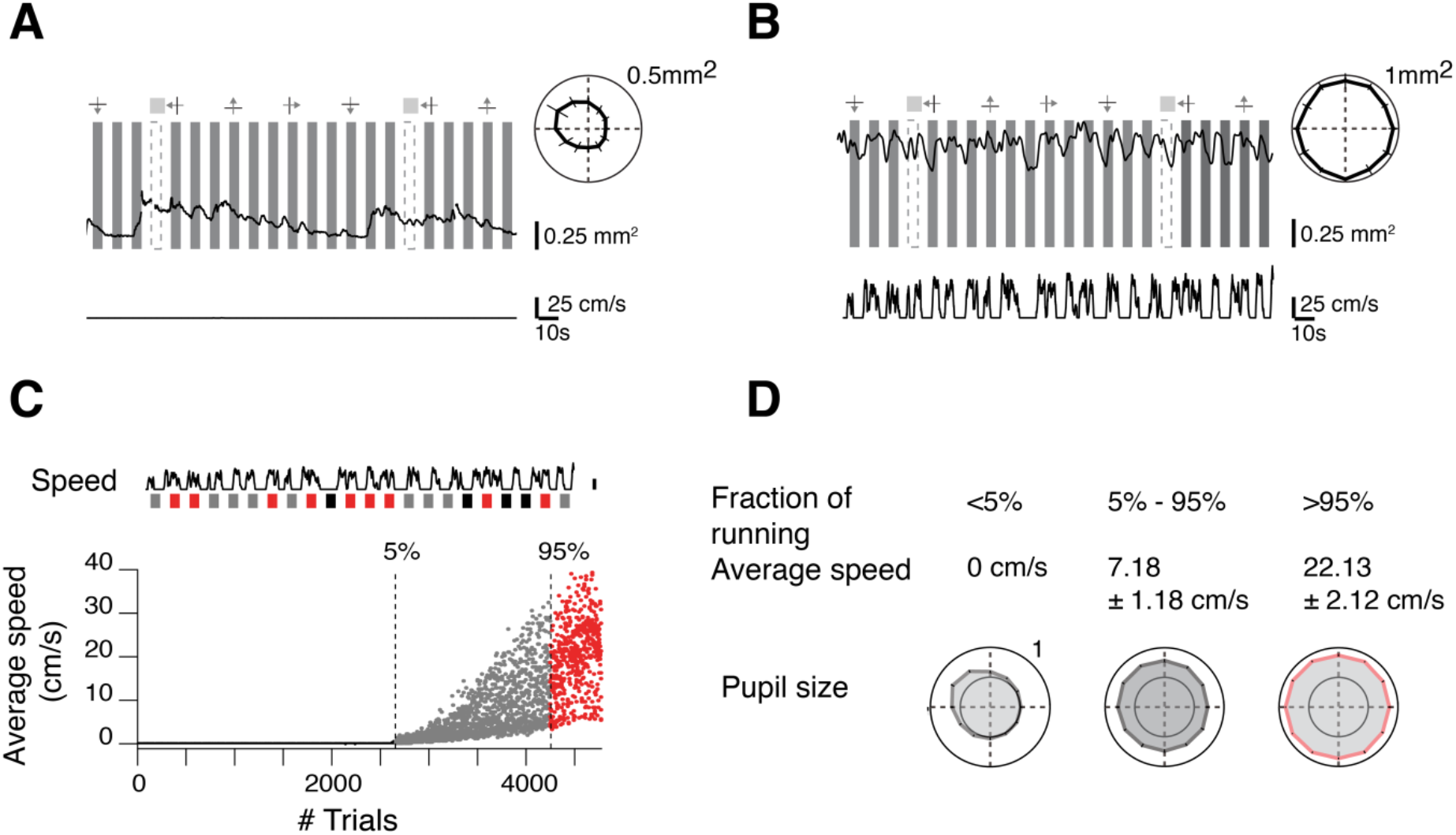
Pupil responses to visual motion reflect a vision-only arousal mechanism. (A) Example pupil response to stimulus direction during stationary and (B) locomoting behaviors. Polar plots represent the average response (10 trials, mean±SD). (C) Selection criteria for behavior. Trials were classified into three groups: stationary (black, <5%running), locomotion (red, >95% running) and mixed (gray, 5%-95% running). Dashed lines represent threshold for stationary and running trials (total number of trials: 4776). In most trials mice did not run (56% stationary trials; 11% locomotion trials). (D) Back-to-front motion stimuli elicit a pupil response during stationary trials. Average pupil size: stationary (25 sessions; included 221±47 trials per stimulus), mixed (15 sessions; 115±43) and locomotion trials (7sessions; 44±9).

Thus, in head-fixed animals, visual stimuli drifting in the back-to-front direction elicit a specific increase in pupil size that does not occur for other stimuli. Consistent with a change in arousal, the effects occurred in absence of locomotion, were strongest when the animal is still and nearly absent when engaged in continuous locomotion.

### Stimulus-induced arousal modulates the amplitudes of dLGN responses

To examine the impact of behavioral state on early visual responses, we measured in awake animals the cellular Ca^2+^ responses of dLGN axons arborizing through the layers of the visual cortex (Figure 3A).We used adeno-associated viral vector injections to label dLGN neurons with the genetically-encoded calcium indicator GCaMP6f (AAV1.CAG.GCaMP6f.WPRE.SV40). Post-mortem confocal imaging of immunohistochemically stained fixed tissue confirmed that somatic GCaMP6 expression was confined to the dLGN, and that GCaMP6f expressing axonal projections were visible in all layers of V1 and were densest in layer 4 (data not shown), as reported previously (Kondo and Ohki, 2015; Sun et al., 2015). Using a video-rate two-photon microscope, we imaged axonal Ca^2+^ responses to the drifting gratings stimuli of different orientations and directions (depth between 20μm – 400μm) (Figure 3B). To study axonal responses to the stimuli, regions of interest (putative presynaptic boutons) were identified by thresholding the maximal projection during visual stimulation and applying morphological operations to outline putative varicosities (Figure 3C). We examined the time course of fluorescence changes (dF/F) of 8260 presynaptic putative boutons (n=14 mice, 37 sessions) in awake state. On average, in each session number of active boutons was 275±145 boutons per imaging plane. We observed boutons responding to different orientations and directions of moving gratings (Figure 3D, Figure S2A, Figure S3A).

**Figure 3.**
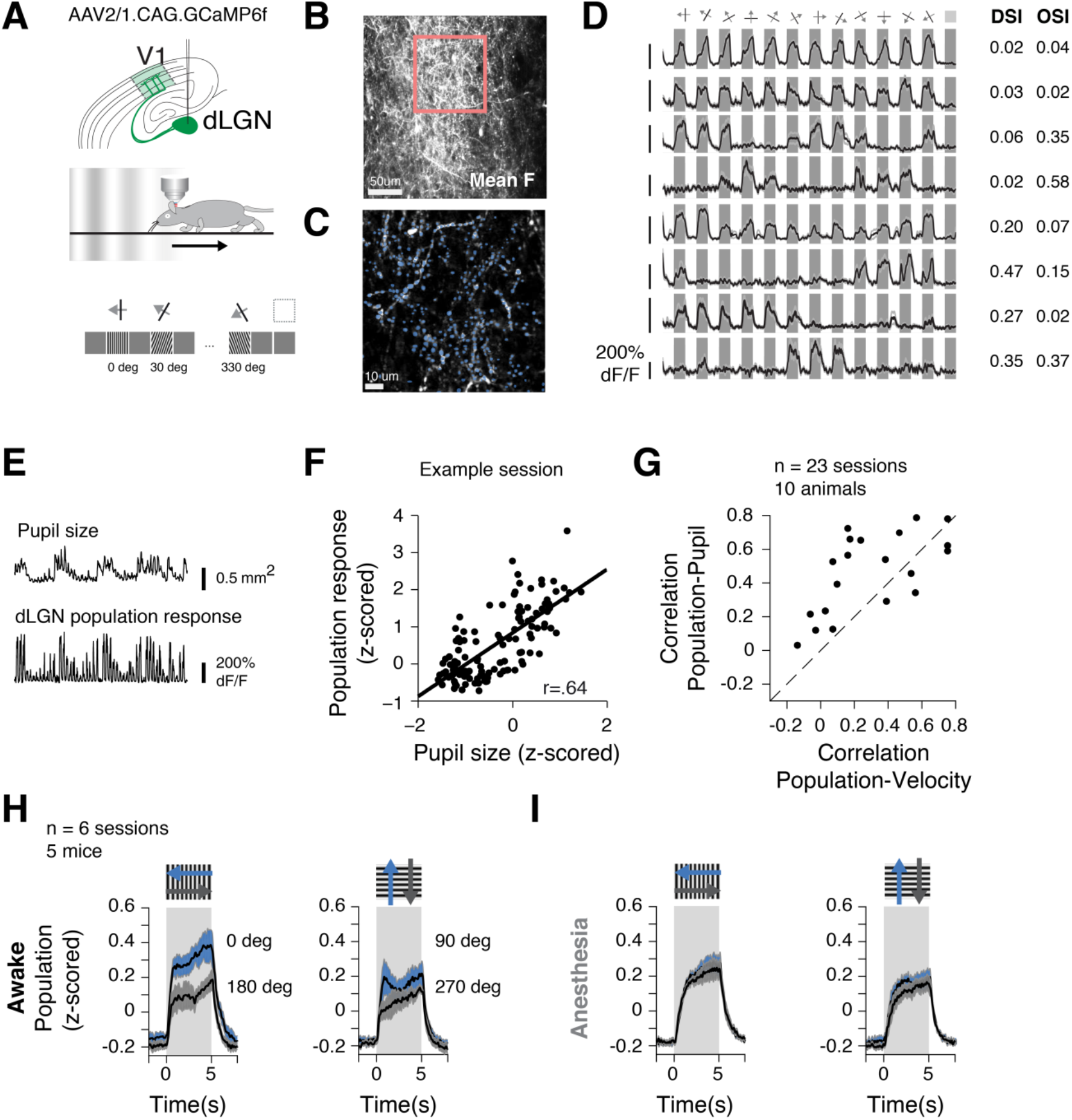
Stimulus-induced arousal biases the amplitude of dLGN responses. (A)Non-invasive method for functional imaging of dLGN axons projecting to the mouse V1 during the assay. Illustration of the strategy used for expressing GCaMP6f in axons (top), experimental setup (middle) and visual stimulus sequence (bottom). (B) Example of an average imaging plane with axonal projections labeled with GCaMP6f in layer 4 of V1 in an awake mouse. (C) Segmentation of putative presynaptic boutons. (D) Average calcium transients in response to gratings drifting in 12 directions. Individual dLGN boutons prefer diverse directions and orientations (average over 10 trials, mean±SD); left: direction- and orientation-selective indices (DSI and OSI). (E-F) population and pupil responses are correlated during visual stimulation; dashed line in F is the linear fit. (G) Correlation between dLGN population responses and animal speed for individual sessions. (H) Average calcium transients of dLGN population for the cardinal directions for the datasets where acquisition preceded measurements under anesthesia (mean±S.E.M). Like pupil responses (see Figure 1F), the amplitude of dLGN responses increases during stimulation in temporal-to-nasal direction. (I) Response amplitudes for the same imaging plane as (H) under anesthesia. The amplitude of responses is comparable in all directions. See also Figure S2.

dLGN population activity was correlated with changes in pupil size, a proxy for arousal state (Figure 3E). We used subset of data accompanied with pupil diameter measurement’s (n=10 mice, 23 sessions). To estimate population activity, we computed the average normalized activity across visually-responsive boutons and then related the resulting activity across time courses to pupil size, speed and visual stimulation. The amplitude of dLGN axon Ca^2+^ responses to visual stimuli were strongly correlated with pupil (Figure 3F, example sessions, Spearman’s rank correlation = 0.64). In most sessions, population response was positively correlated with pupil (0.51±0.25, mean±SD) and with animal speed (0.34±0.30) (Figure 3G).

Stimulus-induced changes in behavioral state modulated the amplitude of population activity. We compared dLGN response to stimuli of different orientation drifting in opposite directions in subset of data preceding measurements under anesthesia (Figure 3H; Figure S3A; n=5 mice, 6 sessions, 1816 boutons). As seen with the pupil size measurements (Figure 1E; Figure S1B), we observed a strong asymmetry in dLGN responses to the visual stimuli of different direction of motion (Figure 3H; Figure S2C). Responses to stimuli moving the back-to-front, temporal-to-nasal directions (Figure 3H; Figure S2C, blue) showed larger amplitudes than stimuli moving in the opposite directions (Figure 3H; Figure S2C, gray).

To demonstrate a causal role of behavioral influences in these neural activity biases, responses measurements in awake animals were immediately followed by measurements under light anesthesia, ensuring response measurements of individual boutons from precisely the same regions of the visual cortex (Figure 3I; Figure S3B; n=5 mice, 6 sessions, 1014 boutons). Measurements in awake and anesthetized had comparable quality (Figure S2A-B; Figure S3A-B;). We found that anesthesia abolished the population-level bias of responses in favor of back-to-front motion relative to front-to-back motion (Figure 3I, Figure S2D, blue and gray lines respectively). We conclude that amplitudes of dLGN Ca^2+^ responses to visual motion stimuli are impacted by the animal’s behavioral response to the stimuli.

### Stimulus-induced arousal biases direction tuning measurements

We next investigated the impact on sensory tuning. Axon imaging studies have reported different direction tuning in awake and anesthetized animals (Kondo and Ohki, 2015; Sun et al., 2015), reporting in awake animals an excess of units tuned to the back-to-front direction not seen under anesthesia. The increased visual response during stimulation with back-to-front stimuli possible explains these differences. To examine the orientation and direction preferences of the responses of our axon data sets, we computed orientation and direction selectivity indices and preferences (Figure 3D; Figure S2A-B) and compared results in awake and anesthetized states.

Consistent with earlier work (Cruz-Martín et al., 2014; Kondo and Ohki, 2015; Roth et al., 2015; Sun et al., 2015), we observed that dLGN axons encode diversely tuned signals for orientation and direction (Figure 4). During wakefulness, nearly a third of dLGN boutons showed tuned responses for direction (Figure 4A, 2487 of 8260 boutons; 30% boutons with DSI> 0.2; DSI=0.13±0.15, median±interquartile range) (Figure S3C, Figure S4A) and orientation (Figure 4D, 4177 of 8260 boutons; 51% boutons with OSI>0.2; OSI=0.20±0.19) (Figure S3E, Figure S4E). Under anesthesia, the fraction of direction tuned decreased significantly (Figure S3D; Figure S4B; 137 of 1014 boutons; 14% boutons; DSI=0.09±0.09,p<10^-10^, Wilcoxon rank-sum test). However the fraction of boutons tuned for orientation was similar (Figure S3F; Figure S4F; 497 of 1014 boutons; 49% boutons; OSI=0.20±0.19, p=0.29, Wilcoxon rank-sum test).

**Figure 4.**
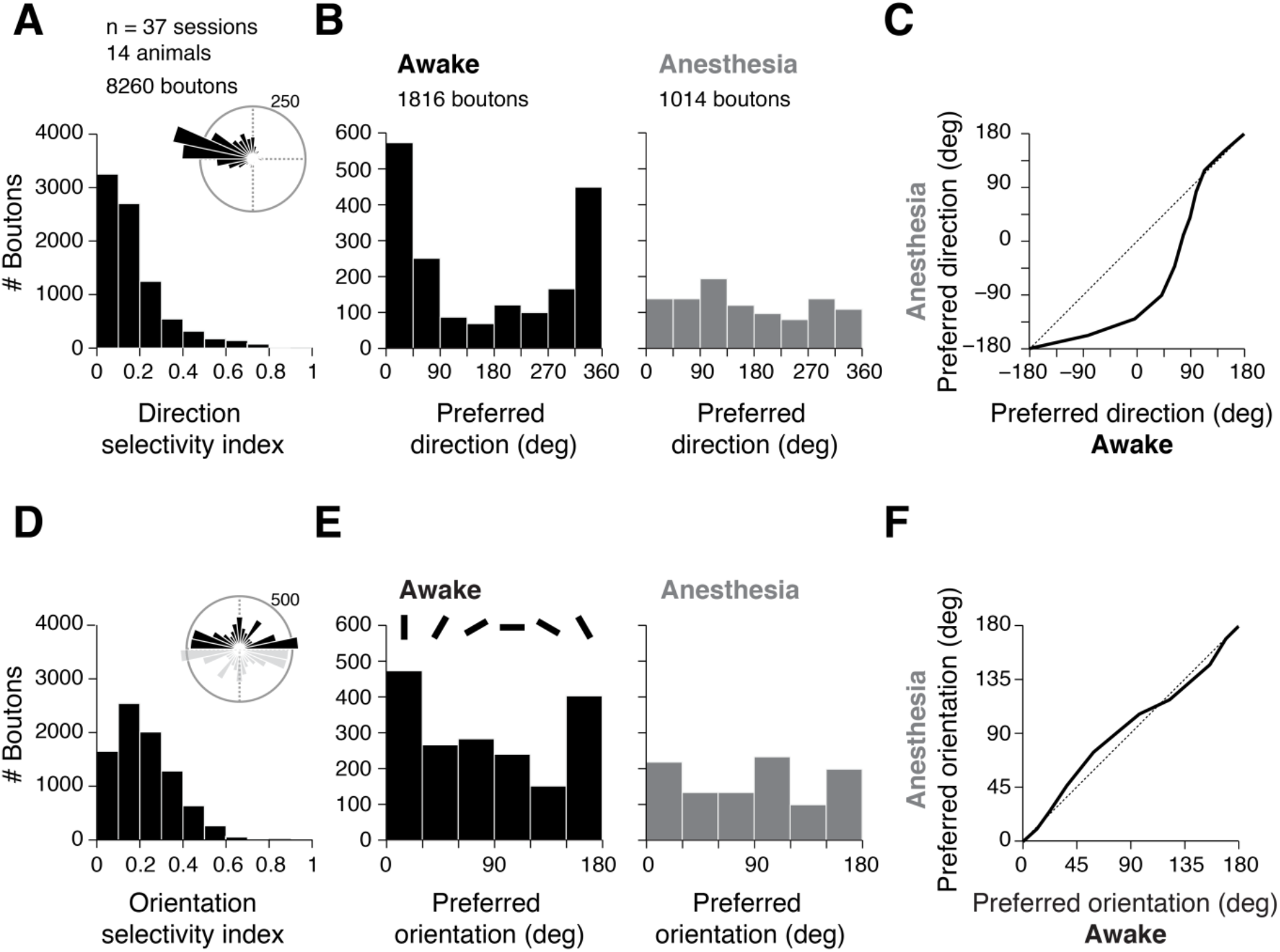
Stimulus-induced arousal biases direction tuning measurements. (A) Distribution of DSI of dLGN boutons in V1. Top right: preferred directions of tuned boutons. (B) Distribution of the preferred directions of all boutons in the subset of sessions in which awake and anesthetized data were acquired from the same imaging planes. We observed the same bias in preferred directions, without thresholding for direction tuning; in anesthesia responses are distributed uniformly in all directions. (C) Quantile-quantile plot of the distributions of preferred directions show differences in awake and anesthesia. Nasal direction is shifted to the center of the plot (0 deg); -180 deg represents temporal direction; directions change clockwise. Diagonal line represents identical distributions. Responses in awake mice are different from anesthesia and biased to the temporal-to-nasal direction. (D) Distribution of OSI of dLGN boutons (same boutons as in (A)). Top right: preferred orientations of tuned boutons. (E) Distribution of the preferred orientation for the same boutons as in (B); bars represent the orientation of gratings. (F) Quantile-quantile plot of the distributions of preferred orientations show minor differences between awake and anesthesia. 90 deg: vertically oriented gratings (nasal-temporal motion directions), 0 deg: horizontally oriented gratings (upward-downward motion directions). See also Figure S3, S4.

During wakefulness, we also observed an excess of boutons tuned to the temporal-to-nasal, back-to-front direction (Figure 4A, inset). This bias was not observed under anesthesia (Figure 4B-C, Figure S4C-D; p= 0.06, Wilcoxon rank-sum test). In contrast, the distributions of orientation preferences were similar in the awake and anesthetized states (Figure 4E-F, Figure S4G-H, p=0.03, Wilcoxon rank-sum test).

To assess the contribution of behavioral modulations to direction tuning of responses, we computed for each imaged bouton a population coupling index defined the correlation coefficient between activity time courses of individual boutons and pupil size (Figure 5A-D). The strength of correlation to population activity was highly predictive of the boutons’ selectivity (Figure 5A) and preferences (Figure 5B). Boutons with strong population coupling showed stronger direction selectivity (Figure 5A) and were often tuned to the back-to-front direction of motion (Figure 5B). A clear link between direction selectivity and population coupling was observed in all sessions (Figure 5C). Most boutons’ responses were strongly correlated with populations activity (Figure 5D).

**Figure 5.**
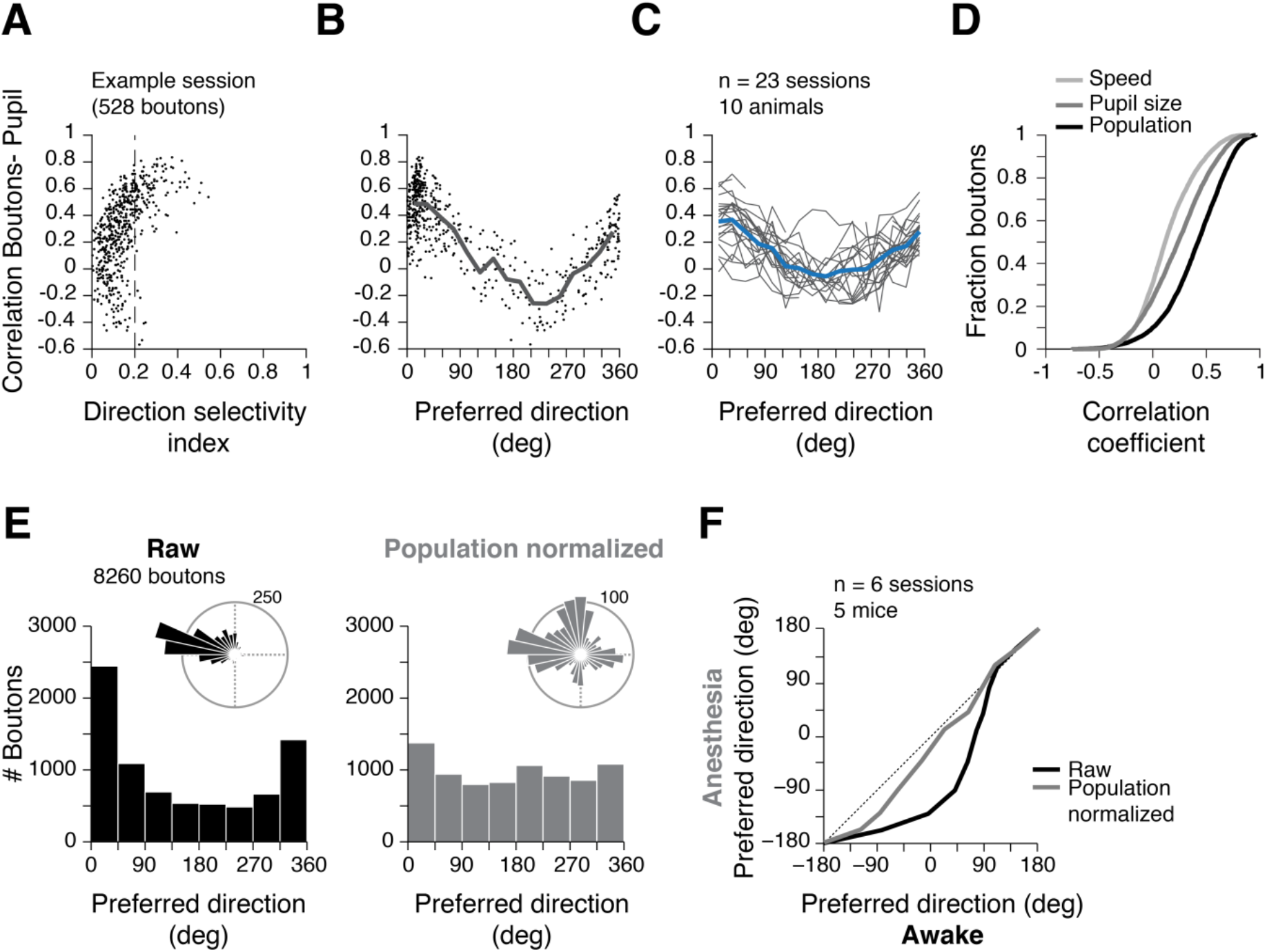
Arousal responses to visual motion impact dLGN population code. (A) Population coupling index quantifying the correlation between activity time courses of individual boutons and pupil. Highly coupled boutons showed stronger direction selectivity indices and (B) tuning preferences (example session). Gray line represents mean value of correlation coefficient calculated to each direction. (C) Same as (B) but for sessions with measurements of pupil, speed and activity time courses; gray lines: individual sessions (lack of continuity in line is due to absence of responses to some directions); blue line: average over all sessions. (D) Cumulative distribution of correlation coefficients between the activity time courses of individual boutons and animal speed (light gray), pupil size (gray) and population response (black). Boutons were highly correlated with population activity. (E) Left: Distribution of preferred direction (same as in Figure 4A). Right: distribution of preferred directions of individual boutons normalized to population response; top: preferred directions of tuned boutons recalculated after normalizing to the population response (23% boutons, DSI>0.2). Population normalization reduced responses biased to the back-to-front direction and revealed responses to other directions. (G) Quantile-quantile plot of the distributions of preferred directions in anesthesia and in awake mice before (same as in Figure 4D, black line) and after normalization (gray line). Differences in distributions become smaller after normalization, approaching the distribution for recordings under anesthesia. See also Figure S5.

To further assess the impact of behavior on dLGN tuning preferences, we normalized the boutons time courses by population activity and computed a corrected set of tuning indices. The impact of normalization on selectivity indexes was modest (Figure S5A-B, raw OSI=0.20±0.19; normalized OSI=0.18±0.18; 45% boutons; p<10^-10^, Wilcoxon rank-sum test; raw DSI=0.13±0.15; normalized DSI=0.12±0.12; 23% boutons; p<10^-10^, Wilcoxon rank-sum test). However, the bias to the back-to-front direction of motion was greatly reduced by normalization (Figure 5E, p<10^-10^, Wilcoxon rank-sum test) and approached that observed in anesthetized animals (Figure 5F, p=0.004, Wilcoxon rank-sum test). Weaker effects were observed on orientation preferences (Figure S5C-D, p=0.04, Wilcoxon rank-sum test). Normalization did not affect direction or orientation preferences of boutons measured under anesthesia (Figure S5E; p= 0.008 (direction), Figure S5F; p=0.3 (orientation) Wilcoxon rank-sum test).

## Discussion

We examined the impact of arousal and behavioral state on neuronal encoding of motion stimuli in the mouse visual thalamus. We found that visual motion stimuli elicit a robust and pronounced arousal-related behavioral response that strongly modulates neuronal responses to the stimuli. The effects on neuronal tuning measurements are profound, strongly biasing population measures of direction selectivity and direction preferences. The results demonstrate that visual motion can exert a potent influence on the behavioral state impacting sensory coding at early stages of visual processing.

Our results indicate that stimulus-driven changes in arousal explain the marked differences in direction selectivity and preferences reported in recent studies. A study in anesthetized animals (Kondo and Ohki, 2015) found a nearly equal representation of the 4 cardinal directions whereas a study in awake animals (Sun et al., 2015) reported a strong bias in favor of the back-to-front, temporal-to-nasal direction. We found pronounced effects in awake animals but not in anesthetized (Figure 4H-I). We also found that the impact on neuronal responses was specific to direction preferences in the temporal-to-basal, back-to-front direction (Figure 5A-C). We showed that normalizing data by population activity equalizes response preferences across the four cardinal directions (Figure 5D-E), which makes dLGN responses more similar to those observed in the retina (Briggman et al., 2011; Kay et al., 2011; Dhande et al., 2013; Yonehara et al., 2013; Sabbah et al., 2017). The response patterns we observed during locomotion, stillness and under anesthesia all point towards stimulus-driven arousal related mechanism.

We showed pronounced effects in dLGN inputs, which provide the main visual input to V1. The effects on pupil size and neuronal responses occur without concomitant locomotor activity which suggests neuromodulators influences. Arousal influences have been described in dLGN (Erisken et al., 2014; Hei et al., 2014) but also in the visual cortex (Reimer et al., 2014, 2016; Zhuang et al., 2014; Vinck et al., 2015). It is therefore possible that similar effects occur in V1. Further investigations are required to determine the impact of these effects on the V1 population code.

The neural circuitry mediating behavioral responses to visual motion is not known. The effects we observe may involve the superior colliculus which, because of its role in attention (Krauzlis et al., 2013), orienting (Sokolov et al., 2002; Stubblefield et al., 2013; Walton et al., 2013) and other innate visual behaviors (Yilmaz and Meister, 2013; De Franceschi et al., 2016; Salay et al., 2018; Shang et al., 2018), is a plausible target. Interestingly, recent studies have shown that the colliculus sends motion information to dLGN (Gale and Murphy, 2014; Bickford et al., 2015; Hillier et al., 2017), which would provide a substrate for the effects.

The function of the arousing influence of back-to-front visual motion is unclear. We observed no concomitant eye movements and locomotor activity that could explain the effects. We also observed no habituation of the behavioral response, which is inconsistent with a startle response. Back-to-front visual motion may particularly salient as it may indicate backward body movement. This could explain why the stimulus was ineffective in inducing pupil size responses during locomotor activity (Figure 2). Experiments in freely moving animals will be required to fully understand the functional significance of the phenomenon described here, particularly during navigation and movement coordination.

## Acknowledgements

We thank Michał Ślęzak for preliminary experiments and analyses that led to this study; Karl Farrow for guidance and comments on the manuscript; João Couto for eye tracking software, helpful discussions, and comments on the manuscript; VB acknowledges support from FWO (Grant G0D0516N), KU Leuven Research Council (Grant C14/16/048) and NERF institutional funding. NERF is funded by Imec, VIB and KU Leuven and the Government of Flanders.

## Author Contributions

KS and VB designed the study; KS performed experiments and analyzed the data with input from VB; MW and DB performed control analyses. KS and VB wrote the paper with input from MW and DB.

## Declaration of Interests

The authors declare no competing interests.

## Methods

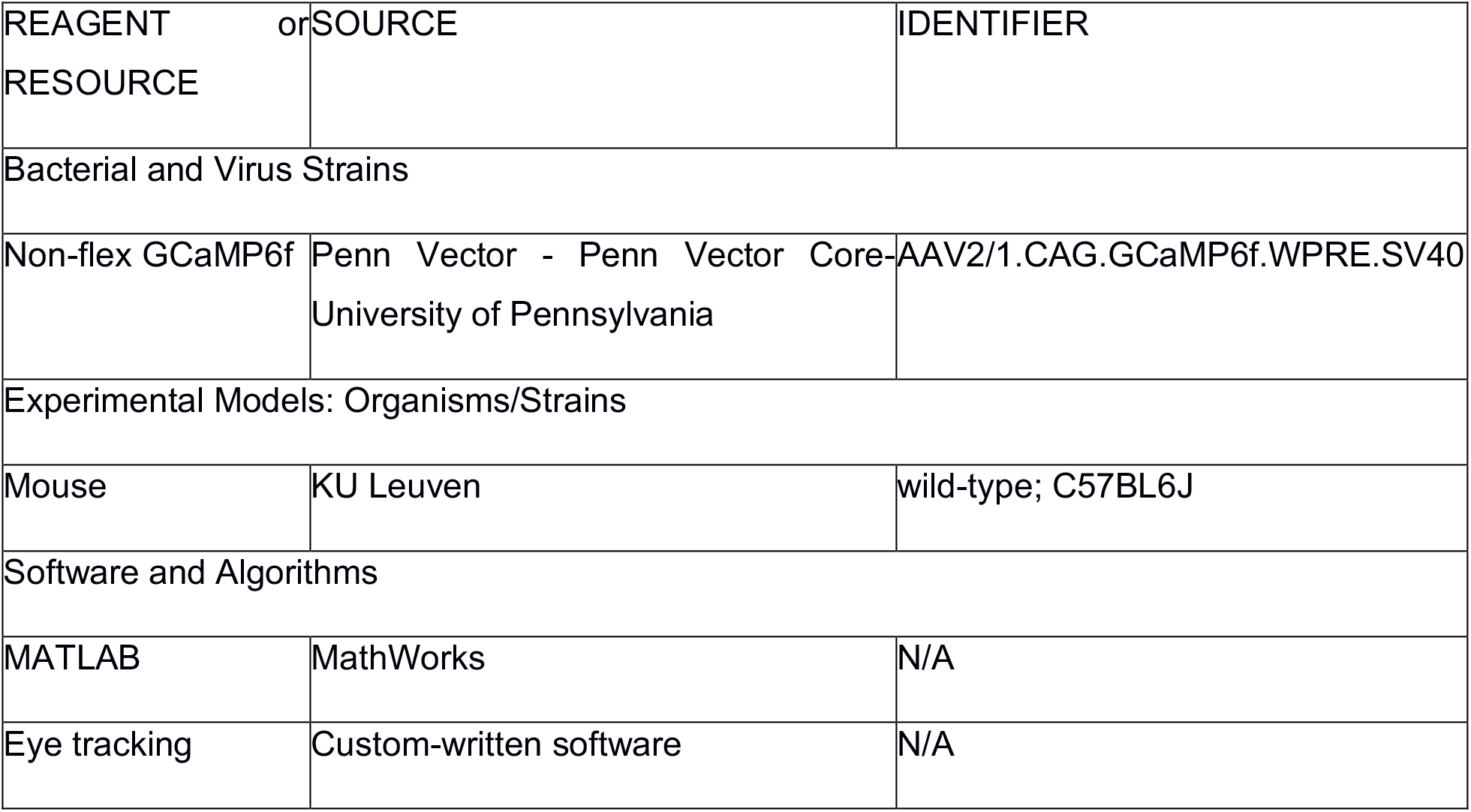
KEY RESOURCES TABLE.

### CONTACT FOR REAGENT AND RESOURCE SHARING

Further information and requests for resources should be directed to and will be fulfilled by the Lead Contact Vincent Bonin

### EXPERIMENTAL MODEL AND SUBJECT DETAILS

#### Mice

All procedures were conducted regarding to ethical protocol approved by the ethical research committee of KU Leuven. We used 18 male mice (wild-type C57BL6J) between 7 and 8 weeks old (23 to 30g) at the day of viral injections. In this study we included 13 mice (pupillometry); 14 mice (calcium imaging); 10 mice (both behavior and calcium imaging); 5 mice (anesthesia). All mice were individually house in the animal facility of NERF.

## METHOD DETAILS

### Surgery and viral injection

Expression of the calcium indicator in thalamic axons was achieved by injecting an adeno-associated virus construct containing GCaMP6f under the CAG promoter (AAV2/1.CAG.GCaMP6f.WPRE.SV40 – University of Pennsylvania Vector Core). Dextamethasone (3.2 mg/kg, i.m.) was delivered at least 4 hours before surgery. Stereotactic coordinates were used to target dLGN (2.10-2.30mm posterior and 2.10-2.25mm lateral from bregma). Mice were anesthetized using isoflurane (3% for induction; 1–1.5% for maintaining). An injection capillary (~15–25 μm internal tip diameter) was lowered to a depth of 2.50-2.80 mm to deliver 40-60 nl of viral construct (Nanoject II, Drummond Sci.). Mice were subsequently implanted with a titanium head plate and a 5mm glass window (Goldey et al., 2014) centered on the primary visual cortex (1.3mm anterior from lambda and 3.1 mm lateral from midline) over the posterior left hemisphere. Post-operative treatment was given for 60 hours following surgical procedures (buprenorphine 0.2 mg/kg i.m. and cefazolin 15 mg/kg I.M. in 12 hours intervals). Thalamic axonal boutons were visible 3 weeks after injection.

### Habituation to head fixation and behavioral assay

After recovery (usually 5 – 7 days surgery), animals were put on a water restriction schedule, and habituated to head-fixation on a 150 cm linear treadmill track (Royer et al., 2012). During training sessions, mice were rewarded with tap water (10 μL drop size) every 150 cm. Mice were trained 2-3 times per week for at least 2 weeks. In the third/fourth week after injection, mice went into two-photon imaging experiments. If the body weight dropped by 15% of the weight before restriction, training was interrupted and free access to water granted.

### Visual stimuli

A 22-inch LED monitor (Samsung 2233RZ, 1680 by 1050-pixel resolution, 60 or 120 Hz refresh rate, brightness 100%, contrast 70%) was positioned 18 cm in front of the right eye, covering 120 by 80 degrees of the visual field (0 to 120 deg. central to peripheral and ± 40 deg. lower to upper visual field). Mice head were tilted by about 15 degrees (so that the microscope objective was perpendicular to the brain surface). Presentation software (Neurobehavioral Systems) was used to control visual stimulation and synchronize behavior readout to the frame rate of the microscope. To characterize orientation and direction selectivity of axonal boutons we used sinusoidal drifting gratings at 12 different directions (30 degrees apart - 13 mice, 33 sessions - or 45 degrees apart - 3 mice, 4 sessions – with SF=0.08 cpd and TF=4Hz). Each stimulus was displayed for 5 seconds with 5 second pre-stimulus blank period (gray uniform screen 45 cd/m^2^). Moving gratings in each direction were displayed in sequence and repeated 5–15 times.

### Locomotion and pupillometry

An optical encoder (Avago Tech) was attached to the shaft of the front treadmill wheel, and used to monitored the position on the track (resolution of ~3 mm). A custom circuit board with a microcontroller (AT89LP52, Atmel) was used to deliver reward after 150 cm. Encoder and reward signals were sampled at 10 kHz by the visual stimulation software using a data acquisition board (MCC, Measurement Computing).

The right eye was illuminated with a 730 nm LED (Thorlabs) and recorded at 30 Hz (AVT Prosilica GC660 with Navitar Zoom 6000; equipped with an infrared filter). Frames were recorded with StreamPix (Norpix) and pupil diameter was extracted offline using mptracker (https://bitbucket.org/jpcouto/mptracker). For each dataset, the eye corners and parameters for morphological operations were manually chosen. Camera frames were smoothed with a 2D Gaussian filter and converted into binary images. Contours were then extracted and fitted with an ellipse. The diameter was computed as the square root of the product of the ellipse axis and converted to mm using an average eye diameter of 6 mm. Position was corrected by the corneal reflection (when present) and converted to spherical coordinates. Traces of pupil diameter were re-sampled to the frame rate of the microscope.

### Two-photon calcium imaging

A custom-built two-photon microscope (Neurolabware) was used to image calcium signals of putative axonal presynaptic boutons within V1 (20 to 380 μm depth, frame rate ~15 fps (1024 lines by 1024 pixel) or ~31 fps (512 lines by 512 pixel) 350 by 280 μm field of view). Boutons were sampled from 20–380μm from the pia (9 sessions at depth 20–70μm, 9 sessions at 140-240 μm, 19 sessions at 310–380μm). GCaMP6f was excited at 920 nm using a MaiTai DeepSee laser and green light was collected by GaAsP photomultiplier tubes (Hamamatsu) through a 16x lens (NA = 0.8, Nikon). Laser power at the objective was 40–145 mW.

For experiments under anesthesia, mice were kept on the linear treadmill and anesthetized with isoflurane (0.5–1 %, 0.5 L/min O2) to keep a similar imaging plane. Eyes were protected with a thin layer of mineral oil. We report data from 6 sessions (5 mice) acquired at a depth 310–360 μm.

## QUANTIFICATION AND STATISTICAL ANALYSIS

All data analysis was performed in MATLAB (The Mathworks).

## Behaviour analysis

The trial selection criteria was modified from (Erisken et al., 2014). A trial was considered during locomotion when the animal was running 95% of the trial at least at 1cm/s. Stationary trials were defined as those were the animal velocity was below 1 cm/s in at least 95% of the trial. 56% of the trials were classified as stationary, 11% locomoting and 33% were not classified (mixed trials).

Pupil size traces were smoothed with a median filter and either normalized to the *95^th^* percentile or z-scored.

## Calcium data analysis

Registration of two-photon imaging frames was done using TurboReg (Thévenaz et al., 1998) by aligning each frame to a target image (generated by averaging a motion-free stretch of 128 frames).Regions of interest (ROI) were identified by thresholding the maximal projection of all frames and applying morphological filtering to outline varicosities. In total 15358 ROI’s were outlined (37 sessions, 14 mice). The time course of calcium responses was calculated as the fractional change *dF/F* = (*F_corrected_* – *F*_0_)/*F*_0_ × 100.0 where *F_0_* is baseline fluorescence and *F_corrected_* = *F_raw_* – *F_motion_* – *F_neuropil_* summing pixel intensities over the bouton’s ROI, and subtracting the estimated contributions of motion and neuropil as in (Bonin et al., 2011).

A bouton was considered as visually-responsive if the responsiveness index was above 1.8. This index was calculated as follows:
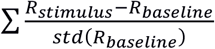,where R_baseline_ is the mean response 2 seconds before the stimulus and *R_stimulus_* is the mean response during the stimulation period.

Orientation tuning was measured using the global orientation selectivity index (OSI):

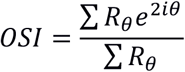

where the response Rθ is the average response across all trials for the direction θ.

Similarly, direction tuning was measured using the global direction selectivity index (DSI), defined as:

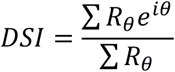

We considered orientation selective bouton if the OSI was above 0.2 and direction selective if the DSI was above 0.2.

Population response of dLGN boutons was calculated by averaging *ΔF/F* of all visual responsive boutons. To correlate dLGN activity with pupil diameter and animal speed, we calculated the Spearman correlation coefficient of z-scored average responses for each session.

Normalization of individual bouton responses was done by dividing the average response during stimulus presentation by the population response. OSI, DSI and stimulus preferences were computed anew.

## DATA AND SOFTWARE AVAILABILITY

Data are available from the corresponding author upon request.

---

**Figure S1.**
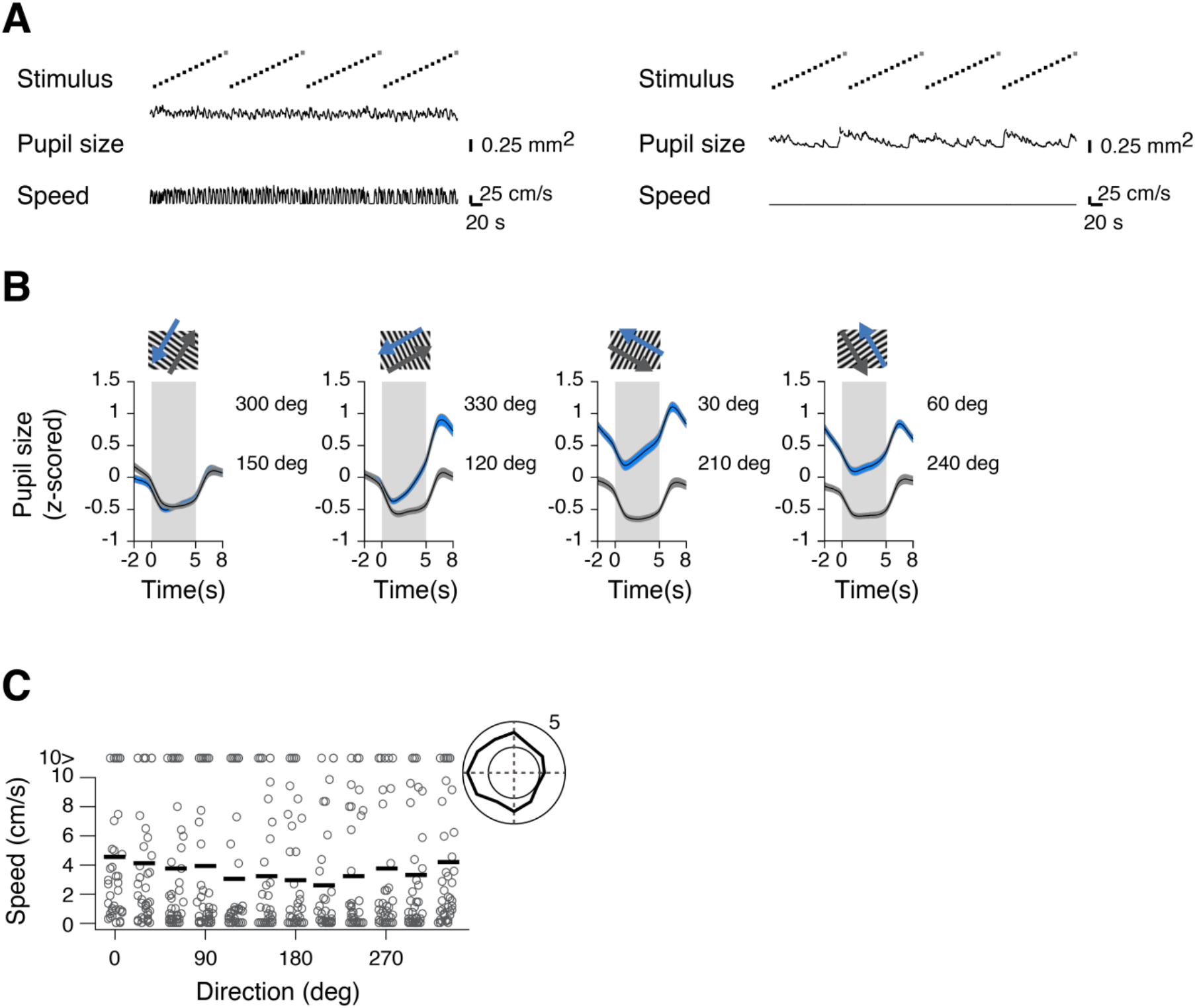
Influence of full-field moving gratings on pupil size and locomotion speed. (A) Example of pupil area and animal speed in distinct mice during presentation of moving gratings to contralateral eye. Left: no effect of stimulus on behavior Right: stimulus influenced arousal with no locomotor response; (B) Normalized pupil responses triggered to the onset of the stimulus for other directions (n=13 mice in 40 sessions). Shaded area is stimulus duration; colors differentiate opposing directions. Pupil increases in response to back-to-front directions: 30 deg and 330 deg. (C) Effect of stimulus direction on animal speed, average over all sessions. Grey dots are average speed for individual sessions during the stimulation epoch. Black lines: mean across all sessions. Polar plot represents the average animal speed in response to moving gratings. On average, animal speed increased during stimulation with gratings moving from the back-to-front.

**Figure S2.**
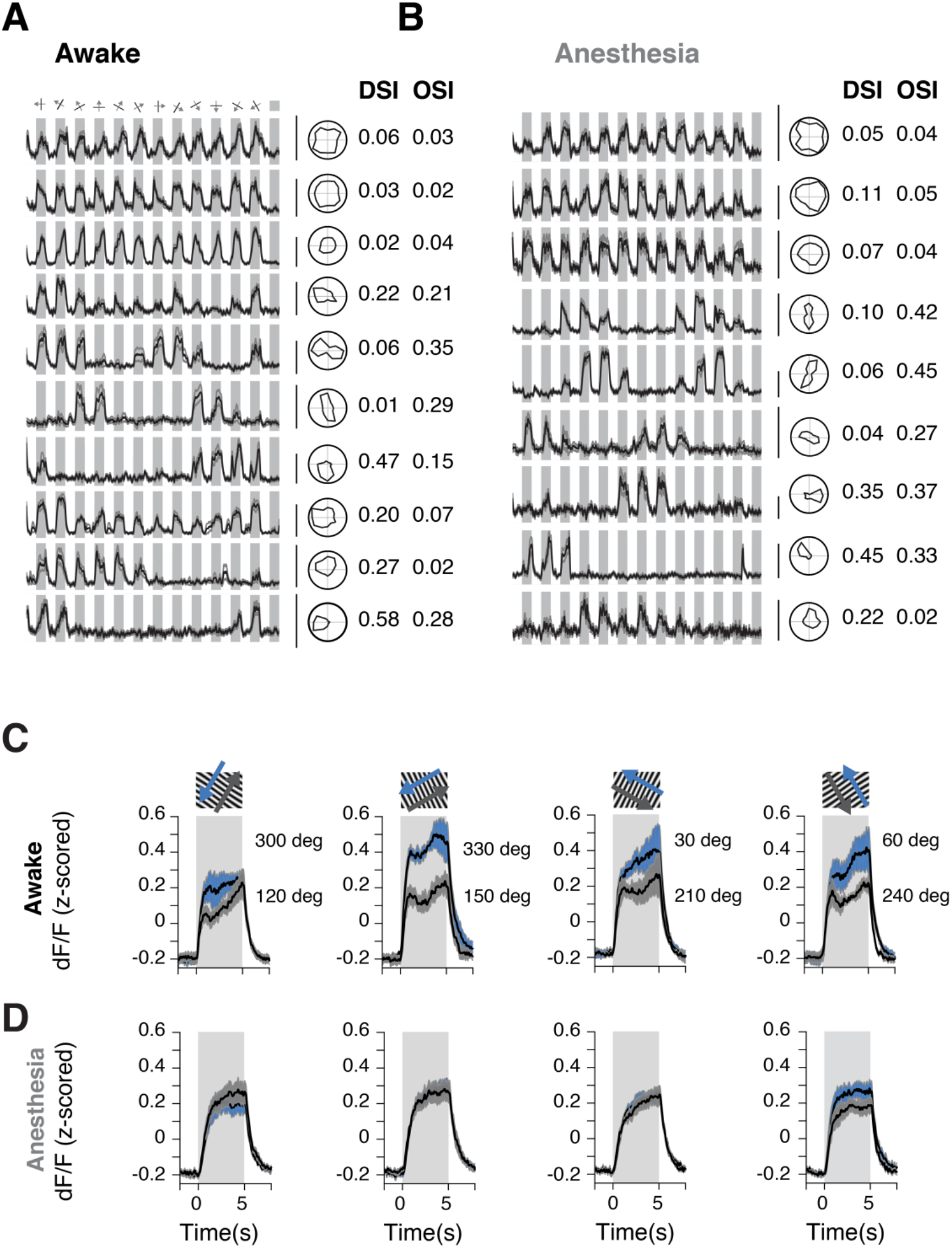
dLGN boutons’ responses in awake and anesthesia. (A) Average calcium transients in response to gratings drifting in 12 directions of boutons in awake sessions (10 trials, average). Middle: polar plots representing tuning properties for bouton; left: OSI and DSI. Examples of bouton responses come from a subset of sessions that preceded measurements under anesthesia (n=5 mice, 6 sessions). Scale bar 100% dF/F. (B) Same as in (A) for anesthesia. Likewise to measurements in the awake state, boutons showed non-selective, direction-selective and orientation-selective responses (10 trials, average). (C) Average calcium transients of dLGN population for other directions (n=5 mice in 6 sessions, mean ± SEM). Colors differentiate contrary directions. (D) Same as in (C) for anesthetized mice (same imaging plane as in (C)).

**Figure S3.**
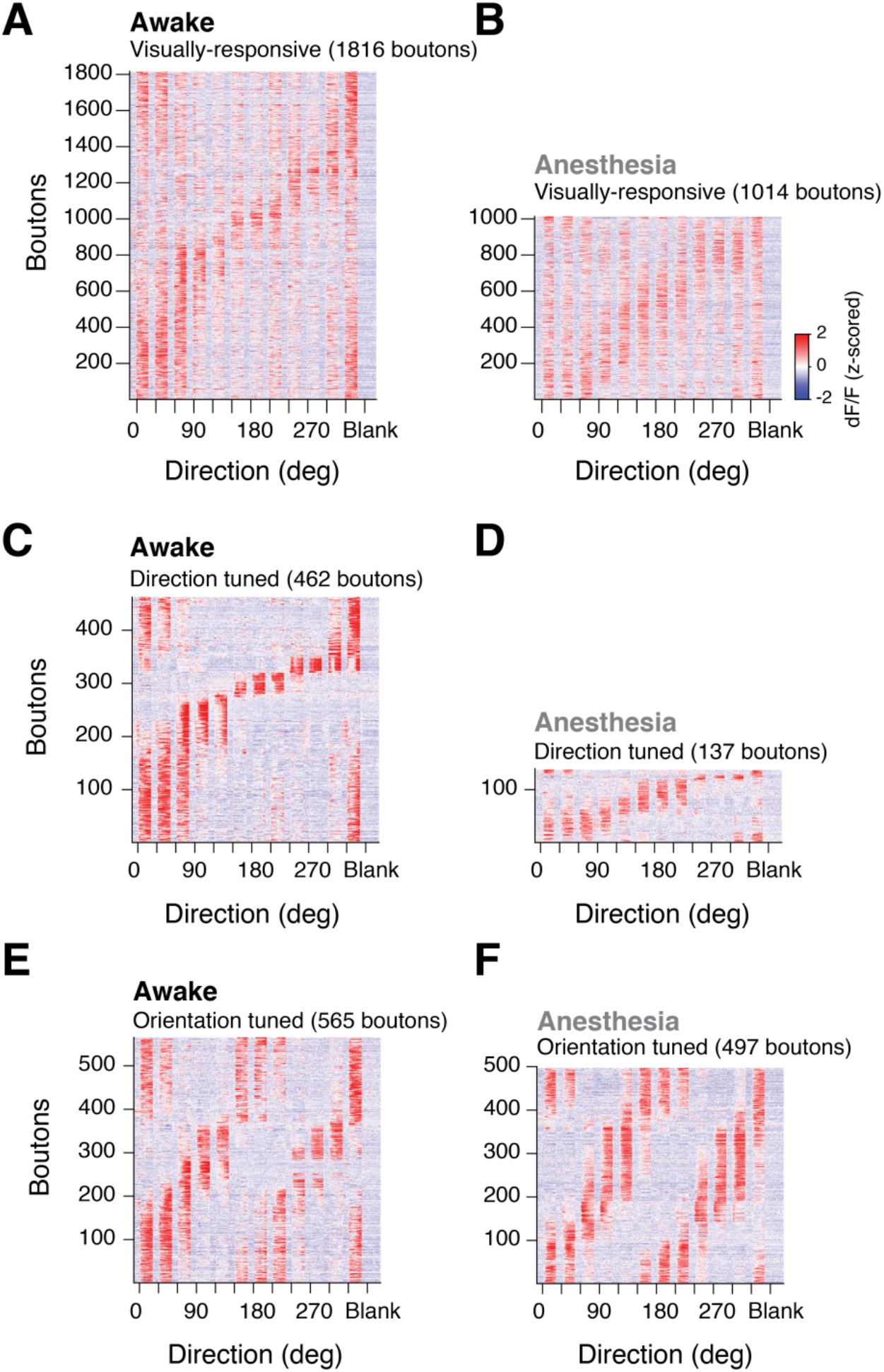
Pronounced reduction of direction-tuned responses under anesthesia. (A) Heat maps of average response of all visually-evoked boutons in awake (left) and (B) anesthesia (right) sessions (same imaging planes). Responses are preceded by 2s of gray screen before stimulus onset. Boutons are sorted by preferred direction. (C-D) Orientation tuned responses (OSI>0.2). (F-G) Direction tuned responses (DSI>0.2).

**Figure S4.**
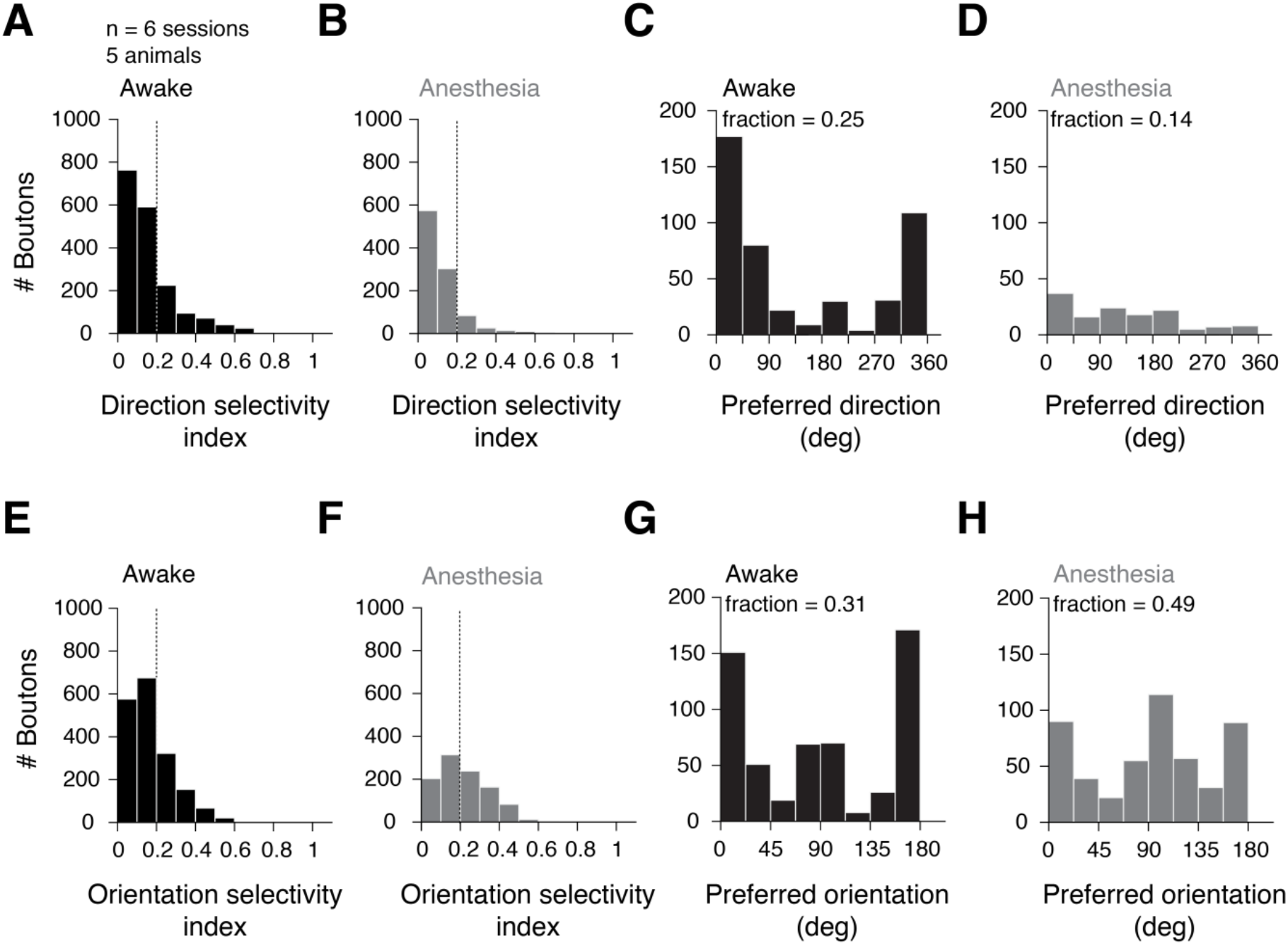
Distributions of DSI and OSI in awake and anesthesia. (A) Distribution of DSI of bouton responses in awake and (B) anesthesia sessions. Dashed line is threshold for tuned responses. (C) Preferred directions of tuned boutons in awake (25% boutons) and (D) anesthesia sessions (14% boutons). (E) Distribution of OSI for dLGN boutons in awake and (F) anesthetized sessions. (G) Preferred orientations of tuned boutons in awake (31% boutons) and (H) anesthesia (49% boutons).

**Figure S5.**
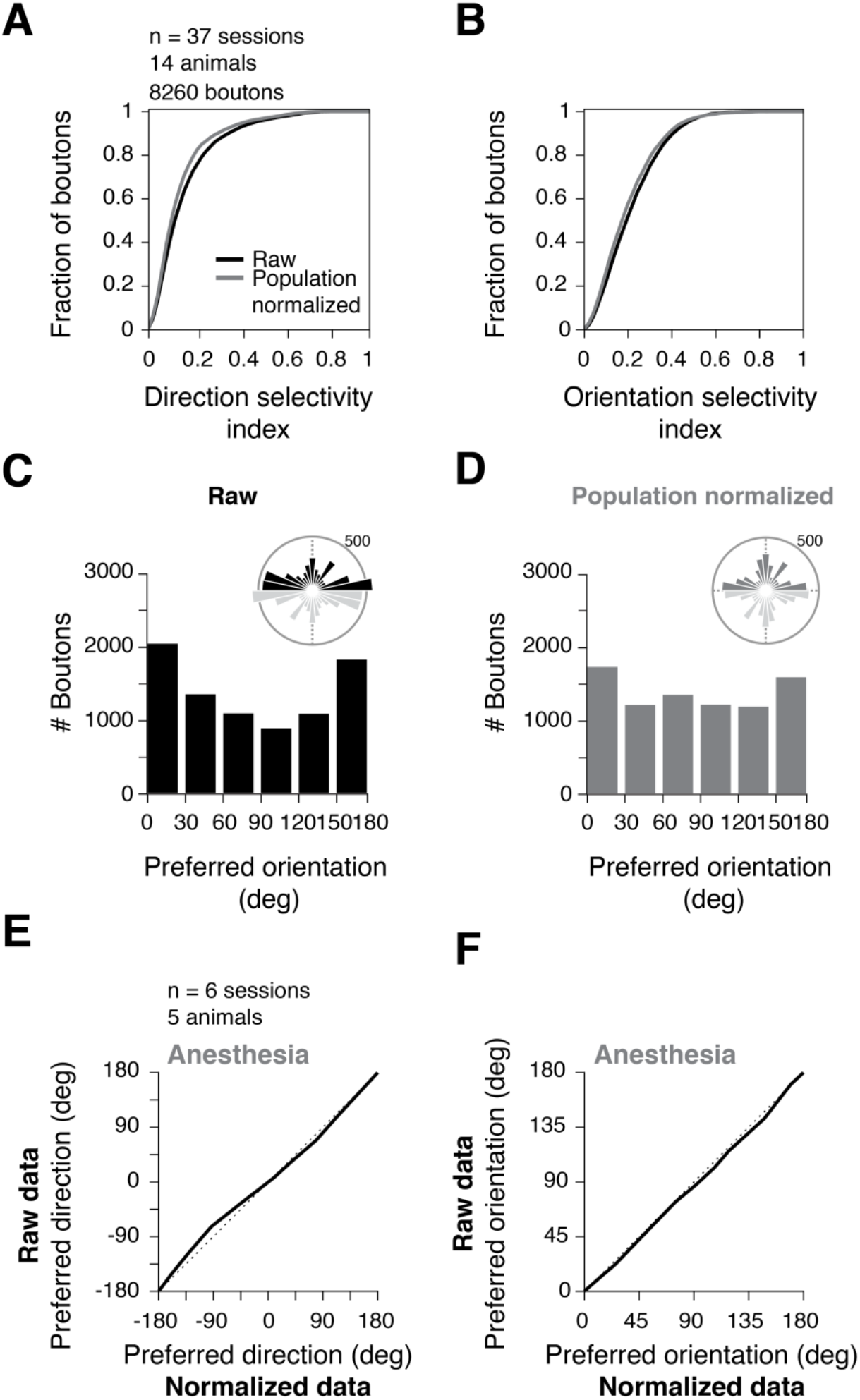
Effect of normalization on dLGN boutons’ tuning properties. (A) Cumulative of the distribution DSIs before (black) and after normalization to population response (gray). DSIs shifted towards lower values. (B) Same as (A) for OSIs. (C) Distribution of preferred orientation of all visually-evoked boutons in 37 sessions; top: preferred orientation for tuned boutons (51% boutons). (D) Same as in (C) recalculated after normalization; top: preferred orientation for tuned boutons (45% boutons). (E) Quantile-quantile plot of the distributions of preferred directions and (F) preferred orientations in anesthetized mice before and after normalization to population response (all boutons). Normalization to population response does not affect data acquired under anesthesia.

